# Center of mass states render multi-joint torques throughout standing balance recovery

**DOI:** 10.1101/2024.08.14.607976

**Authors:** Kristen L. Jakubowski, Giovanni Martino, Owen N. Beck, Gregory S. Sawicki, Lena H. Ting

## Abstract

Successful reactive balance control requires coordinated modulation of hip, knee, and ankle torques. Stabilizing joint torques arise from feedforward neural signals that modulate the musculoskeletal system’s intrinsic mechanical properties, namely muscle short-range stiffness, and neural feedback pathways that activate muscles in response to sensory input. Although feedforward and feedback pathways are known to modulate the torque at each joint, the role of each pathway to the balance-correcting response across joints is poorly understood. Since the feedforward and feedback torque responses act at different delays following perturbations to balance, we modified the sensorimotor response model (SRM), previously used to analyze the muscle activation response to perturbations, to consist of parallel feedback loops with different delays. Each loop within the model is driven by the same information, center of mass (CoM) kinematics, but each loop has an independent delay. We evaluated if a parallel loop SRM could decompose the reactive torques into the feedforward and feedback contributions during balance-correcting responses to backward support surface translations at four magnitudes. The SRM accurately reconstructed reactive joint torques at the hip, knee, and ankle, across all perturbation magnitudes (R^2^>0.84 & VAF>0.83). Moreover, the hip and knee exhibited feedforward and feedback components, while the ankle only exhibited feedback components. The lack of a feedforward component at the ankle may occur because the compliance of the Achilles tendon attenuates muscle short-range stiffness. Our model may provide a framework for evaluating changes in the feedforward and feedback contributions to balance that occur due to aging, injury, or disease.

**NEWS AND NOTEWORTHY:** Reactive balance control requires coordination of neurally-mediated feedforward and feedback pathways to generate stabilizing joint torques at the hip, knee, and ankle. Using a sensorimotor response model, we decomposed reactive joint torques into feedforward and feedback contributions based on delays relative to center of mass kinematics. Responses across joints were driven by the same signals, but contributions from feedforward versus feedback pathways differed, likely due to differences in musculotendon properties between proximal and distal muscles.

## BACKGROUND

When responding to postural perturbations during standing, individuals rapidly produce corrective torques that are coordinated across the hip, knee, and ankle, arising from both feedforward and feedback pathways (Fig 1) (1). The fastest the nervous system can generate a corrective torque through neurally-mediated sensory feedback pathways is approximately 100 ms—which includes both the conduction time and also the neuromechanical delay (2, 3). During this delay, the reactive torque to perturbations arises from the properties of activated muscle where short-range stiffness is tuned by anticipatory feedforward muscle activation (4, 5). Simulations suggest that the torque generated through anticipatory feedforward control can influence the torque generated via feedback control (6, 7). Moreover, aging, injury, or disease can alter both feedforward and feedback pathways, impacting balance control in these individuals. For example, older adults exhibit longer feedback delays (2, 3), and increased feedforward muscle activation (8), where the feedforward muscle coactivation is thought to be a compensatory mechanism for increased feedback delays (6). Differentiating the feedforward and feedback contributions to the overall response would further the fundamental understanding of balance control and may also help identify specific mechanisms underlying balance impairments for the development of targeted rehabilitation. By leveraging knowledge about the delays associated with feedforward and feedback control, here, we sought to identify the feedforward and feedback contributions to balance-correcting joint torque responses at the hip, knee, and ankle.

**Figure 1.**
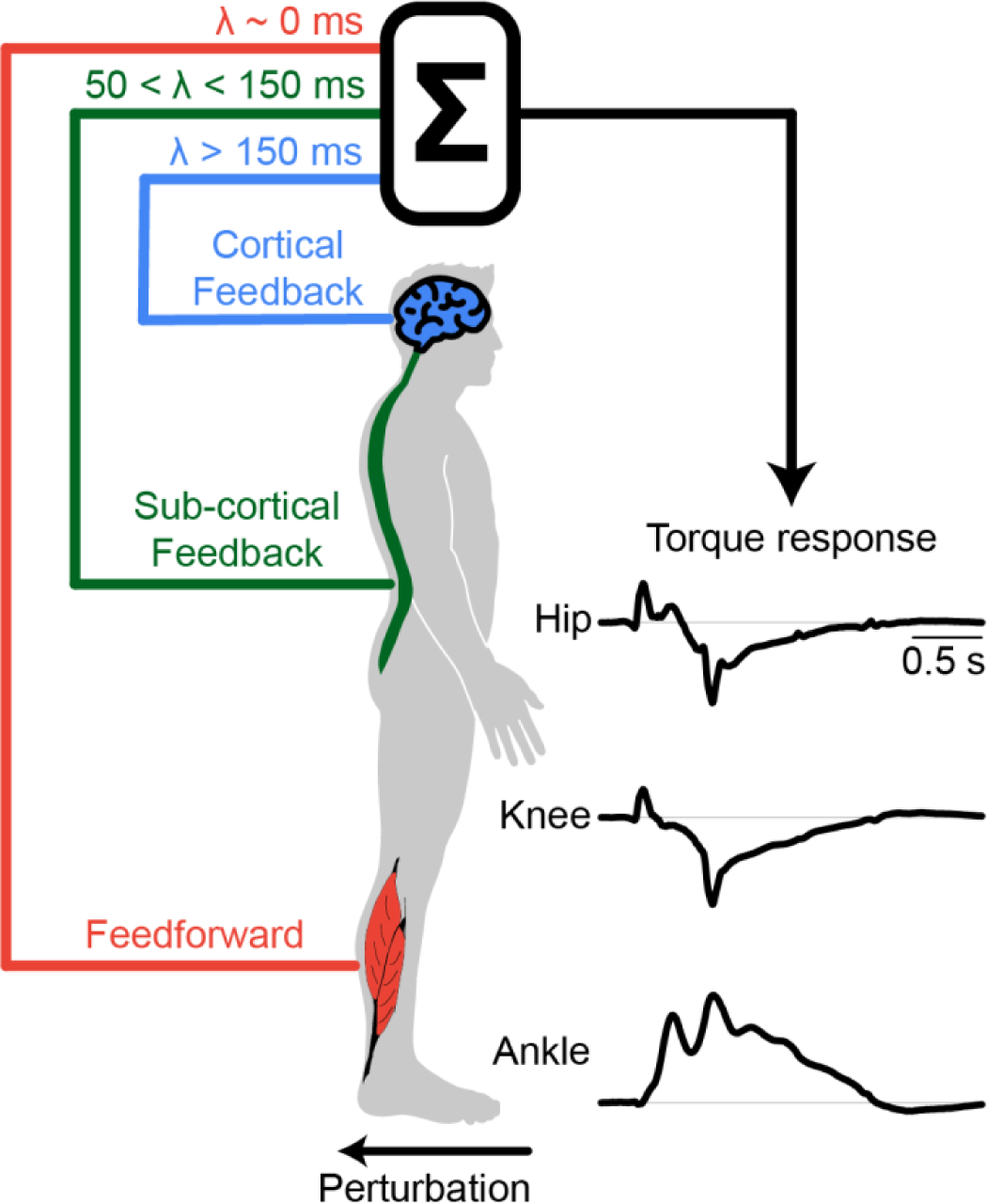
Schematic of the balance correcting response. The torque response to postural perturbations at each joint is mediated by the neurally-mediated feedforward pathways, where the torque produced at the time of the perturbation (λ ∼ 0 ms), as well as neurally-mediated sub-cortical (50 < λ < 150 ms) and cortical (λ > 150 ms) pathways.

Feedforward balance control includes the muscle activation patterns prior to the perturbation that modulate muscle short-range stiffness, providing instantaneous resistance to unexpected perturbations (9, 10). Human standing posture is inherently unstable. Thus, for a delayed feedback system—which the human nervous system is in part—to stabilize the body, the time constant of the body has to be larger than the inherent delay of the feedback sensorimotor system (11), with larger time constants increasing the available time for the nervous system to react within a safety margin of the feedback response. The nervous system can modulate the time constant of body instability through feedforward muscle activation and co-contraction (12). The force generated via muscle short-range stiffness is modulated by the nervous system by the muscle activation level prior to the perturbation (5, 13), while co-contraction increases the short-range stiffness of multiple muscles spanning the joint, further increasing the resistance to unexpected perturbations (14). Modeling studies suggest that the nervous system may leverage feedforward muscle co-contraction during postural control in the presence of noise as a means to minimize the energetic cost compared to solely relying on feedback control (14). Additionally, individuals increase feedforward muscle co-contraction to increase postural stiffness when balance is challenged or threatened (15-19). However, feedforward strategies alone are insufficient to stabilize the body; thus, feedback control is also required to maintain postural stability (20-22).

Feedback responses are generated through sensorimotor transformations, where the nervous system receives sensory information and translates it into reactive muscle activations that generate joint torque. The delay between the onset of a perturbation (e.g., the change in sensory feedback) to the onset of joint torque depends upon the sensory feedback pathway (e.g., subcortical or cortical pathways; Fig 1) and the neuromechanical delay—the latency between neural drive and muscle force production. We previously demonstrated that an error-based sensorimotor transformation of the delayed center of mass (CoM) kinematics (e.g., acceleration, velocity, and displacement) robustly explains reactive muscle activations (23). The sensorimotor response model (SRM) is based on the principle that the neuromuscular system coordinates the activation of muscles across the body to maintain task-level goals, such that coordinated muscle activations reflect task-relevant errors (e.g., CoM displacement) as opposed to joint-level errors (24). We have extensively used the SRM to predict feedback muscle activations across multiple joints and different perturbation conditions (23-27). Most recently, the EMG-SRM, through the implementation of parallel loops with independent parameters, has dissociated components of the long-latency ankle muscle response from subcortical versus cortical pathways (28). However, muscle intrinsic torque responses that arise due to neurally mediated feedforward activation of muscles are not accounted for in the EMG response to a perturbation, but could be considered an instantaneous feedback response at the level of joint torque.

One advantage of using a torque-SRM over the previously developed EMG-SRM is the potential to identify the feedforward muscle short-range stiffness response and to dissociate it from the feedback response to postural perturbations. However, it’s unclear if the same physiological principle that underlies the muscle activation response also underlies the multi-joint torque response. Afschrift et al. (29) recently used a modified version of the SRM to estimate the sensorimotor feedback torque response about the ankle during balance recovery during standing and walking. However, it is unclear whether a torque-SRM can predict the response at the hip and knee because prior modeling work suggests that the hip and knee both exhibit a feedforward muscle short-range stiffness response while the ankle does not (30). The SRM is a feedback model; thus, it is possible that it poorly predicts the hip and knee response because of the feedforward, short-range stiffness component in the torque response (30). However, it is also possible that it may be able to capture the short-range stiffness response because the short-range stiffness appears as an instantaneous “biomechanical feedback” response to the perturbation (4, 7, 31).

Here, we evaluated whether a delayed CoM-feedback model could predict the multi-joint torque response to backward support-surface translations during standing. Specifically, we hypothesized that CoM kinematics (acceleration, velocity, displacement) modulate the multi-joint reactive torque response to postural perturbations from both feedforward and feedback neural control mechanisms. Second, we predicted that a torque-SRM could differentiate feedforward and feedback contributions to the torque response at each joint. To test these hypotheses, we examined the reactive torque response at the hip, knee, and ankle to backward support surface perturbations at four different magnitudes. We modified the previously developed multi-loop EMG-SRM to enable the prediction of joint torques (28). The new CoM-driven torque-SRM consisted of four parallel loops, so the input and output of each loop were the same, but each loop had independent gains and delays (See *Sensorimotor Response Model (SRM)*). We demonstrate the utility of a delayed CoM-feedback model for predicting the balance-correcting torque response at the hip, knee, and ankle, and its ability to identify the feedforward and feedback mechanisms contributions to the overall response.

## METHODS

### Participants

Eight healthy young adults (4 females and 4 males; age 25 ± 4 years; height 1.74 ± 0.08 m; mass 71 ± 8 kg) participated in this study. All participants reported no history of neurological or musculoskeletal disorders. The Emory Institutional Review Board approved the study, and all methods were carried out according to the approved protocol (IRB00082414).

### Data collection

This work is part of a larger study, and a portion of the data presented here has previously been published (21). Participants were instructed to maintain balance during ramp-and-hold support surface translations while standing on a custom platform (Factory Automation Systems, Atlanta, GA). Participants stood on force plates embedded in the platform (AMTI, Watertown, MA, USA). Ground reaction forces were collected at 1000 Hz. Participants were instructed to stand with their bare feet 22 cm apart, with their weight evenly distributed between both feet and their arms crossed about their torso. Participants wore a 33-marker set based on a modified version of the Vicon Plug-in Gait model (32) that included additional foot markers (fifth metatarsal, medial and lateral heel, and medial malleolus).

Surface electromyography (EMG) data were collected at 1000 Hz from the medial gastrocnemius, soleus, tibialis anterior, rectus femoris, vastus medialis, biceps femoris, and gluteus medius on the left leg (Motion Lab Systems, Inc., Baton Rouge, LA, USA). Standard skin preparation methods were performed prior to electrode placement (33), and electrodes were placed on the belly of the muscle. Electromyography (EMG) signals were amplified to maximize the signal resolution in each channel. All kinetic and EMG data were synchronized with kinematic data (collected at 100 Hz) using a motion capture system (Vicon, UK, Oxford).

To identify a set of increasingly challenging perturbations for each individual, we first quantified balance capacity by determining each participant’s step threshold to backward support surface translations (i.e., the platform moved the participant’s feet posteriorly). Step threshold was defined as the maximum translation magnitude where participants could maintain balance without taking a corrective step or being caught by the safety harness (21, 34-36). We used an adaptive method running fit (AMRF) algorithm from the Palamedes toolbox (37), which progressively increased (if no step was taken) or decreased (if a step was taken) the magnitude of the platform translation starting at 15cm. For each perturbation, platform acceleration and velocity were scaled with displacement such that braking occurred ∼500 ms after perturbation onset. Catch trials (e.g., forward perturbations) were randomly interspersed to reduce anticipatory motor adaptations (ratio 1 to 4).

Once the step threshold was identified, participants completed 40 ramp-and-hold support surface perturbations set at 12cm and ∼75%, 85%, and 95% of their step threshold (Fig 2, Table 1). To mitigate adaptation and anticipation, participants also experienced 8 cm forward surface translations randomly interspersed within the perturbation set. A 5-minute seated rest break followed every 20 perturbations to mitigate fatigue.

**Figure 2.**
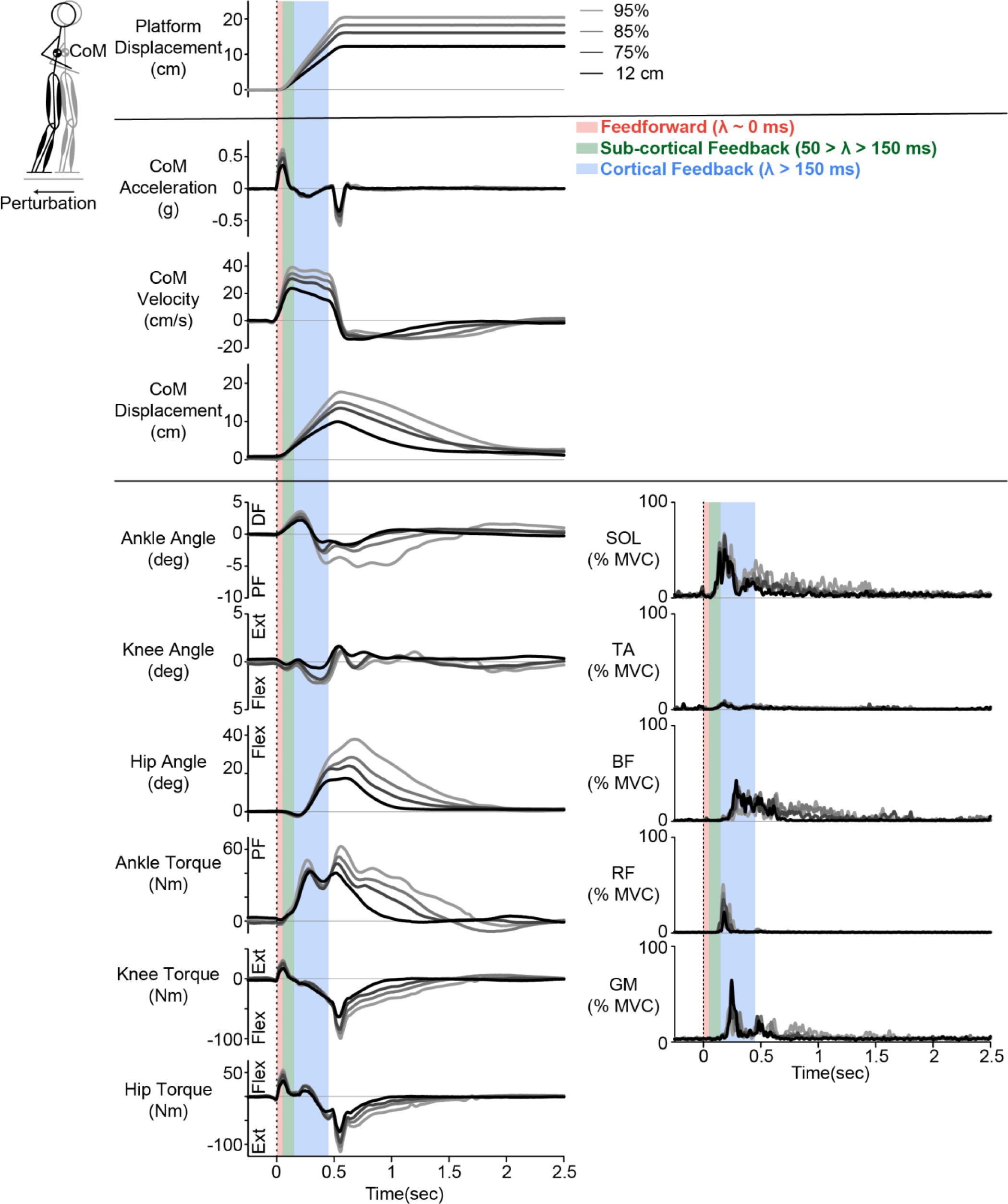
Experimental protocol from a representative participant. Participants were instructed to maintain a foot-in-place balance response to perturbations at 4 magnitudes: 12cm, and 75%, 85%, and 95% of their step threshold. Joint kinematics and kinetics were estimated using the OpenSim Gait 2892 model (38). All torques and angles represent the change in torque from the baseline, pre-perturbation value. The dashed line indicates the start of the perturbation. CoM = center of mass, DF = dorsiflexion, PF = plantarflexion, Ext = extension, Flex = flexion, Sol = soleus, TA = tibialis anterior, BF = biceps femoris, RF = rectus femoris, GM = gluteus medius.

**Table 1:**
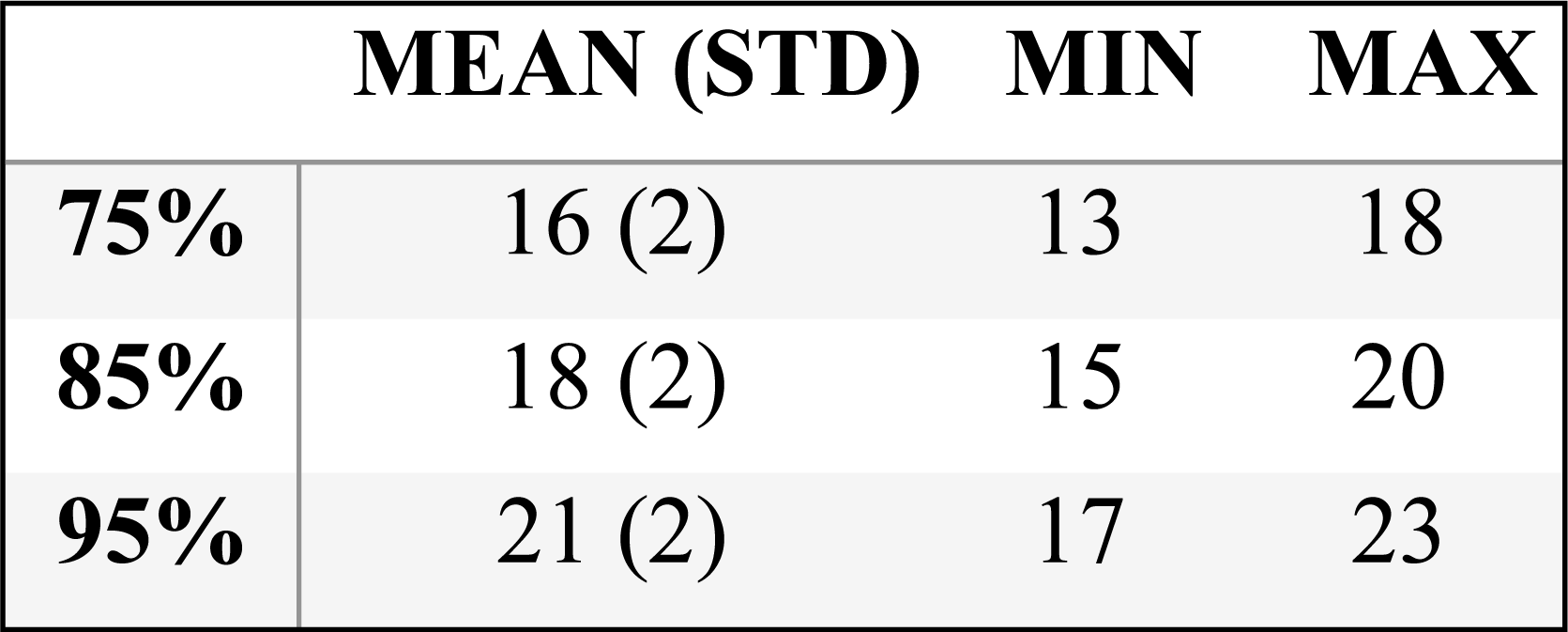
Perturbation magnitudes (cm)

### Data processing

Limb segment marker data and ground reaction forces were used for all estimates of joint kinematics and kinetics. Ground reaction forces were filtered using a fourth-order low-pass filter with a 50 Hz cutoff, while marker data was filtered similarly with a 10 Hz cutoff. Torques at the ankle, knee, and hip were estimated using the inverse dynamics toolbox in OpenSim (Gait 2892 model) (38). We calculated horizontal CoM acceleration as the ground reaction forces divided by the participant’s mass and the platform acceleration, while CoM displacement and velocity were calculated as the weighted sum of the segmental masses from the kinematic data. CoM displacement and velocity were up-sampled using linear interpolation to 1000 Hz for all further analysis.

All EMG data were high-pass filtered using a third-order zero-lag Butterworth filter with a 35 Hz cutoff. They were then demeaned, rectified, and low-pass filtered (40 Hz) (27). EMG signals were then normalized by the peak activity over all analyzed trials, yielding a value between 0 and 100.

Perturbation trials that elicited a stepping response or trials where participants uncrossed their arms were excluded from further analyses. Stepping responses were identified as trials in which the magnitude of ground reaction forces for either leg dropped below 10 N.

### Sensorimotor response model (SRM)

To test our hypothesis that the CoM kinematics modulate the multi-joint reactive torque response to postural perturbations, we modified the previously developed EMG-SRM to reconstruct ankle, knee, and hip torque (25, 27, 31, 32, 39). The previously developed EMG-SRM reconstructs reactive muscle activations as a linear combination of CoM kinematics at a common delay (Eq 1).

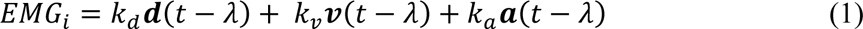

where *k_d_*, *k_v_*, and *k_a_* are the feedback gains on CoM displacement (*d*), velocity (*v*), and acceleration (*a*), and *λ* is the time delay. We note that CoM kinematics represent the deviation of the CoM from a steady-state trajectory relative to the base of support (e.g., the feet), where during standing balance, any change of the CoM resulting from the perturbation is the CoM deviation.

We made two main modifications to this model so it could reconstruct joint torques (Fig 3). The EMG-SRM was developed to examine muscle-level responses, which only contribute in one direction, as muscles can only pull. In contrast, joint torques represent the net effect of the activation of all the muscles that span that joint. This has two implications. First, the torque response (the output) has positive and negative components corresponding to the agonist and antagonist muscle activity. Second, the agonist and antagonist muscle activity is activated differently by the acceleration and braking of the CoM (the input) responses (e.g., agonist muscles are activated in response to CoM acceleration while antagonist muscles respond to CoM braking) (40). Thus, to capture these aspects of the response, parallel loops were added to capture the positive and negative torque response and to predict the torque response to both CoM acceleration and braking (e.g., the positive and negative components of the input; Fig 3). Ultimately, for all joints, the torque-SRM had a maximum of four loops, each with independent gains and delays (Fig 3).

**Figure 3.**
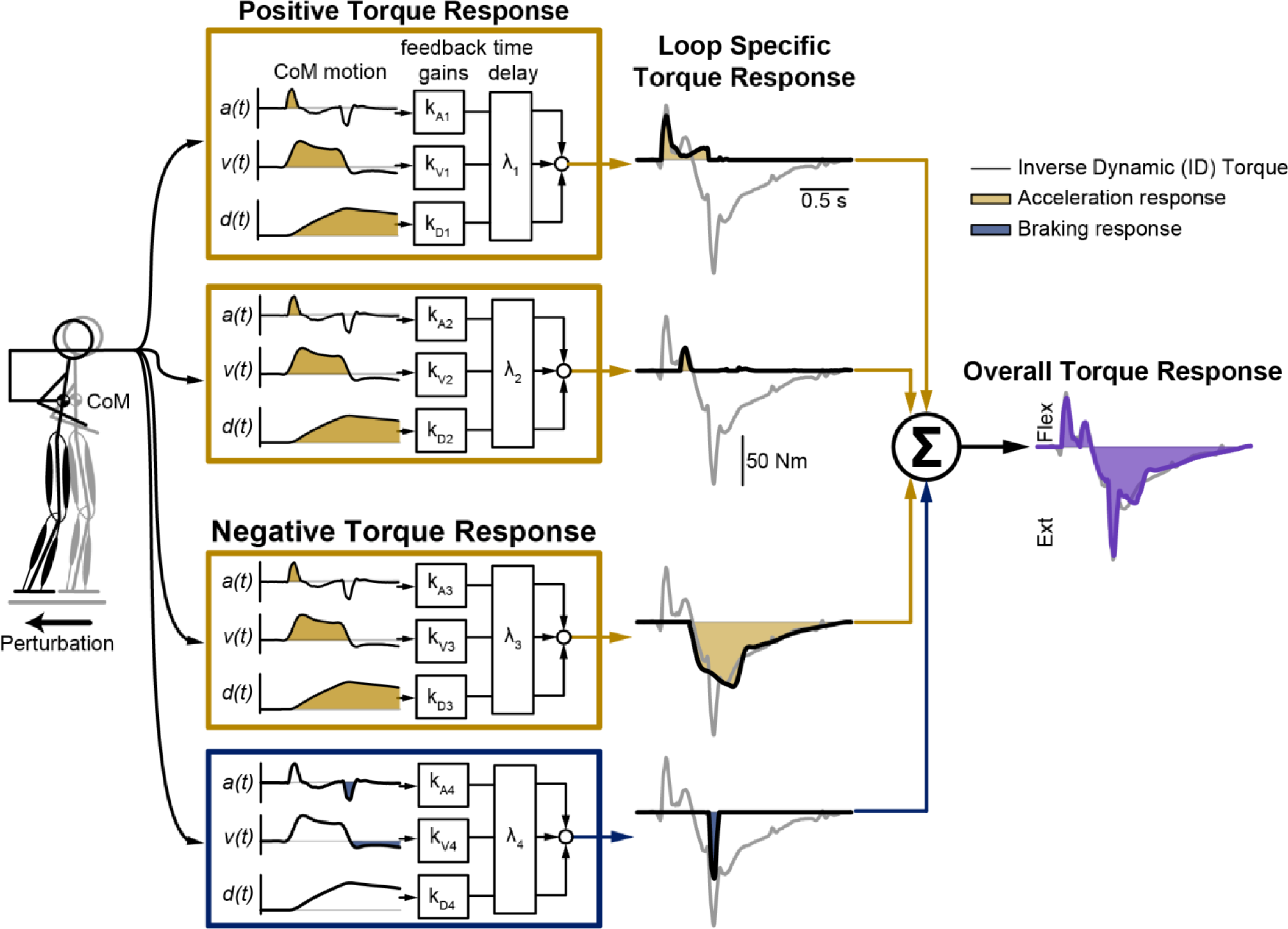
Torque sensorimotor response model (SRM). The SRM predicted joint torque as the quasi-linear sum of CoM deviation (acceleration, a; velocity, v; and displacement, d). We added parallel SRMs, each with independent gains and delays, to predict the torque response. The parallel loops enabled us to predict the positive and negative components of the torque response as well as the torque response to CoM acceleration and braking. Note that this is the model for the hip flexion torque response.

We tuned the gains and delays within each loop to optimize the fit for each participant and perturbation magnitude. All optimizations were performed in Matlab R2022a (Mathworks, Natick, MA) and used the interior point algorithm implemented in *fmincon.m*. First, the trials at the same perturbation magnitude were averaged for use in all further analyses. Next, the background torque was identified as the mean torque one-second proceeding the onset of the perturbation, and this was removed from the overall torque response prior to SRM fitting. For the two SRM loops reconstructing either the positive or negative torque responses, a single, optimization was performed to identify *k_di_*, *k_vi_*, *k_ai,_* and *λ_i_* (where *i* indicates the *i*th SRM loop). Bounds were placed on each loop to prevent the loops from reconstructing the same features within the response (Table 2). Hand-tuning optimization was used to adjust the bounds to achieve the best model fit (e.g., the highest R^2^ and variance accounted for (VAF)). After the two separate optimizations identified the best values of the parameter sets, the gains and delays were concatenated into an initial guess for a final optimization. The final optimization concurrently optimized the gains for both loops with the lower and upper bounds for the gain parameters set at ±10% of the initial optimization, and the bounds for the delay parameters set at ±10ms of the initial optimization. The gains and delays were set to zero if the addition of the loop did not improve the quality of the fit. During the fitting, we found that four loops were required to reconstruct the reactive hip torque, and three loops were required at the knee and ankle (Fig 5, 7 & 9).

**Table 2:**
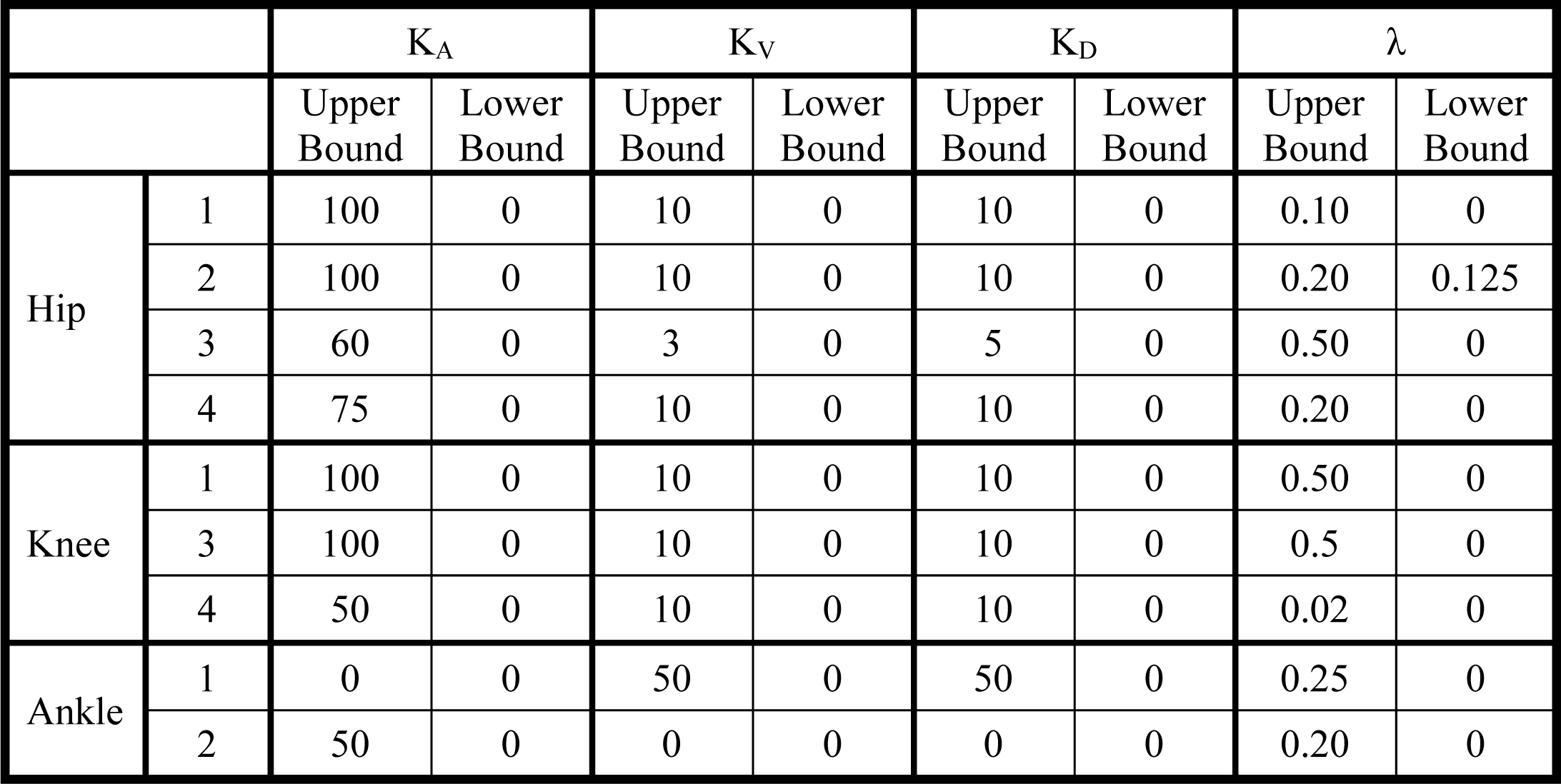
Constraints on sensorimotor response model (SRM) gains.

| | | K<sub>A</sub> | | K<sub>V</sub> | | K<sub>D</sub> | | $\lambda$ | |
| --- | --- | --- | --- | --- | --- | --- | --- | --- | --- |
|  |  | Upper Bound | Lower Bound | Upper Bound | Lower Bound | Upper Bound | Lower Bound | Upper Bound | Lower Bound |
| Hip | 1 | 100 | 0 | 10 | 0 | 10 | 0 | 0.10 | 0 |
|  | 2 | 100 | 0 | 10 | 0 | 10 | 0 | 0.20 | 0.125 |
|  | 3 | 60 | 0 | 3 | 0 | 5 | 0 | 0.50 | 0 |
|  | 4 | 75 | 0 | 10 | 0 | 10 | 0 | 0.20 | 0 |
| Knee | 1 | 100 | 0 | 10 | 0 | 10 | 0 | 0.50 | 0 |
|  | 3 | 100 | 0 | 10 | 0 | 10 | 0 | 0.5 | 0 |
|  | 4 | 50 | 0 | 10 | 0 | 10 | 0 | 0.02 | 0 |
| Ankle | 1 | 0 | 0 | 50 | 0 | 50 | 0 | 0.25 | 0 |
|  | 2 | 50 | 0 | 0 | 0 | 0 | 0 | 0.20 | 0 |

### Statistical analysis

We quantified how well the torque-SRM could reconstruct the reactive joint torques. We quantified the similarity between the inverse dynamics (ID)-derived joint torques and the SRM reconstructed joint torques using R^2^ (squared center Pearson’s correlation coefficient) and VAF. VAF was defined as the square of Pearson’s uncentered correlation coefficient (41) (as has been done in previous studies (27, 39, 40)). We also estimated the root mean square error (RMSE) between the inverse dynamic torque and the SRM reconstructed torque. The RMSE was normalized by the range of the inverse dynamic torque.

We evaluated how the feedback gains changed as a function of perturbation magnitude. We compared the magnitude of the feedback gains and delays using a linear mixed effects model for each joint and each gain or delay. Perturbation magnitude was treated as a fixed factor, while subject was treated as a random factor. For all models, we used a restricted maximum likelihood method to approximate the likelihood of the model and Satterthwaite corrections for degrees of freedom (42). These adjustments reduce Type 1 errors, even for small sample sizes (42). We performed all statistical analyses in MATLAB R2022a. Significance was set *a priori* at α = 0.05. We used Bonferroni post hoc corrections for multiple comparisons. All metrics are reported as the mean ± standard deviation unless otherwise noted.

## RESULTS

### A center of mass-driven sensorimotor response model accurately predicts the reactive multi-joint torque response to perturbations

The SRM qualitatively reconstructed the time history of the torque response at the hip, knee, and ankle at all perturbation magnitudes, capturing the salient features of the response (Fig 4 A, B & C). For example, in the reactive torque response at the hip, we were able to capture both flexion peaks immediately after perturbation onset, as well as the later extension peak (Fig. 4A). The torque-SRM captured peaks in knee and ankle torques as well (Fig 4 B & C). Notably, for all joints, the SRM also captured the torque response after the ramp perturbation ended (>∼0.5 seconds), and could capture all salient features over up to 2.5 seconds after the perturbation (the longest time after the perturbation we could evaluate).

**Figure 4.**
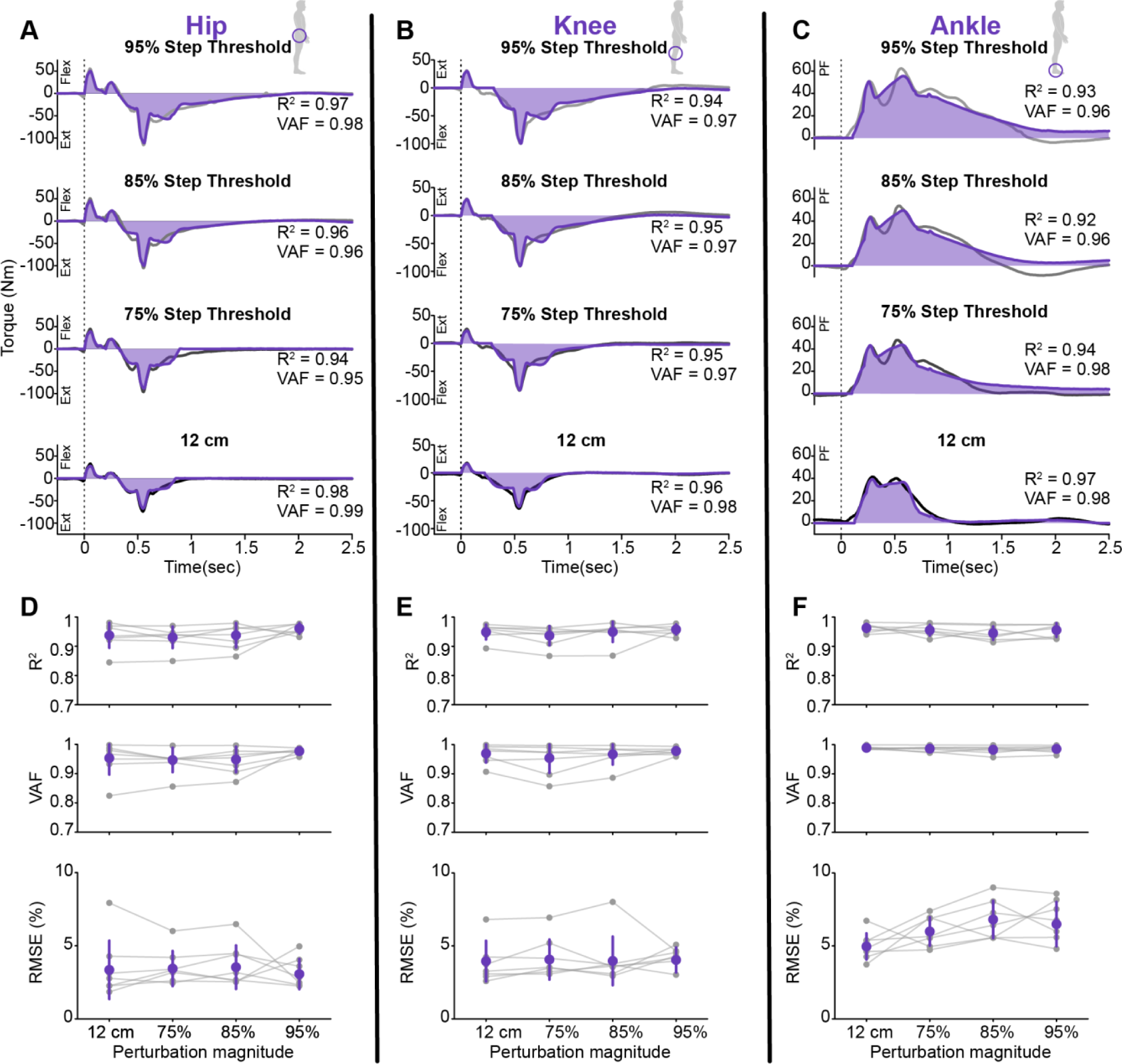
Across all joints, the SRM could accurately reconstruct the torque response at the hip, knee, and ankle. (A - C) Representative fits for the CoM-driven torque-SRM for all perturbation magnitudes (12cm, and 75%, 85%, and 95% of step threshold). The dashed line indicates the start of the perturbation; the SRM fit is in purple, with the inverse dynamic derived torque in black. PF = plantarflexion, Flex = flexion, Ext = extension. (D - F) The SRM reconstructed the inverse dynamics derived torques well at all joints across all perturbation magnitudes. Moreover, there was a low root mean squared error (RMSE) at all joints at all perturbation magnitudes (e.g., <∼10%) between the SRM reconstructed and ID torques. The purple dots represent the group means and standard deviation, while the gray dots and lines represent each participant.

Quantitatively, the SRM accurately predicted the reactive torque response at all perturbation magnitudes at the hip, knee, and ankle (Fig 4 D, E & F). Across all perturbation magnitudes and joints, the SRM was able to accurately reconstruct the reactive joint torques, with high R^2^ (*Ankle:* average: 0.95 ± 0.02, min: 0.91; *Knee:* average: 0.95 ± 0.03, min: 0.87; *Hip:* average: 0.94 ± 0.04, min: 0.84) and VAF (*Ankle:* average: 0.99 ± 0.01, min: 0.96; *Knee:* average: 0.97 ± 0.03, min: 0.86; *Hip:* average: 0.96 ± 0.04, min: 0.82). Moreover, the root mean squared error (RMSE) was low at all joints and magnitudes (*Ankle:* average: 6 ± 1%, max: 9%; *Knee:* average: 4 ± 1%, max: 8%; *Hip:* average: 3 ± 1%, max: 8%).

### A center of mass-driven sensorimotor response model dissociates the feedforward and feedback contributions to the multi-joint reactive torque response

Based on the delays associated with each loop within the SRM, we dissociated feedforward and feedback torque responses, as well as different feedback response pathways at each joint. In this section we discuss differences in the feedforward and feedback contributions at each joint, all driven by task-level feedback of CoM deviations.

**Hip Response:**

Four loops were required at the hip to fit the reactive torque response, with a feedforward loop corresponding to either the acceleration and braking of the CoM, and two feedback loops (Fig 5). The first SRM loop was driven by the acceleration of the CoM at the onset of the ramp perturbation and captured the initial flexion torque at the onset of the perturbation and occurred at nearly a “zero-delay” (average *λ_1_* = 1 ± 1 ms across all perturbation magnitudes) prior to neurally-mediated reactive muscle activation (Fig 2). We thus attribute the zero-delay joint response to feedforward torque arising from the intrinsic properties of hip flexor muscles, including muscle short-range stiffness that is elicited due to the initial extension of the hip (Fig 2). This initial torque response from the hip flexors is counter to the required balance-correcting response to a backward support surface perturbation. Because the CoM is pushed forward, a reactive torque response from the posterior chain muscles is required to maintain balance (43).

**Figure 5.**
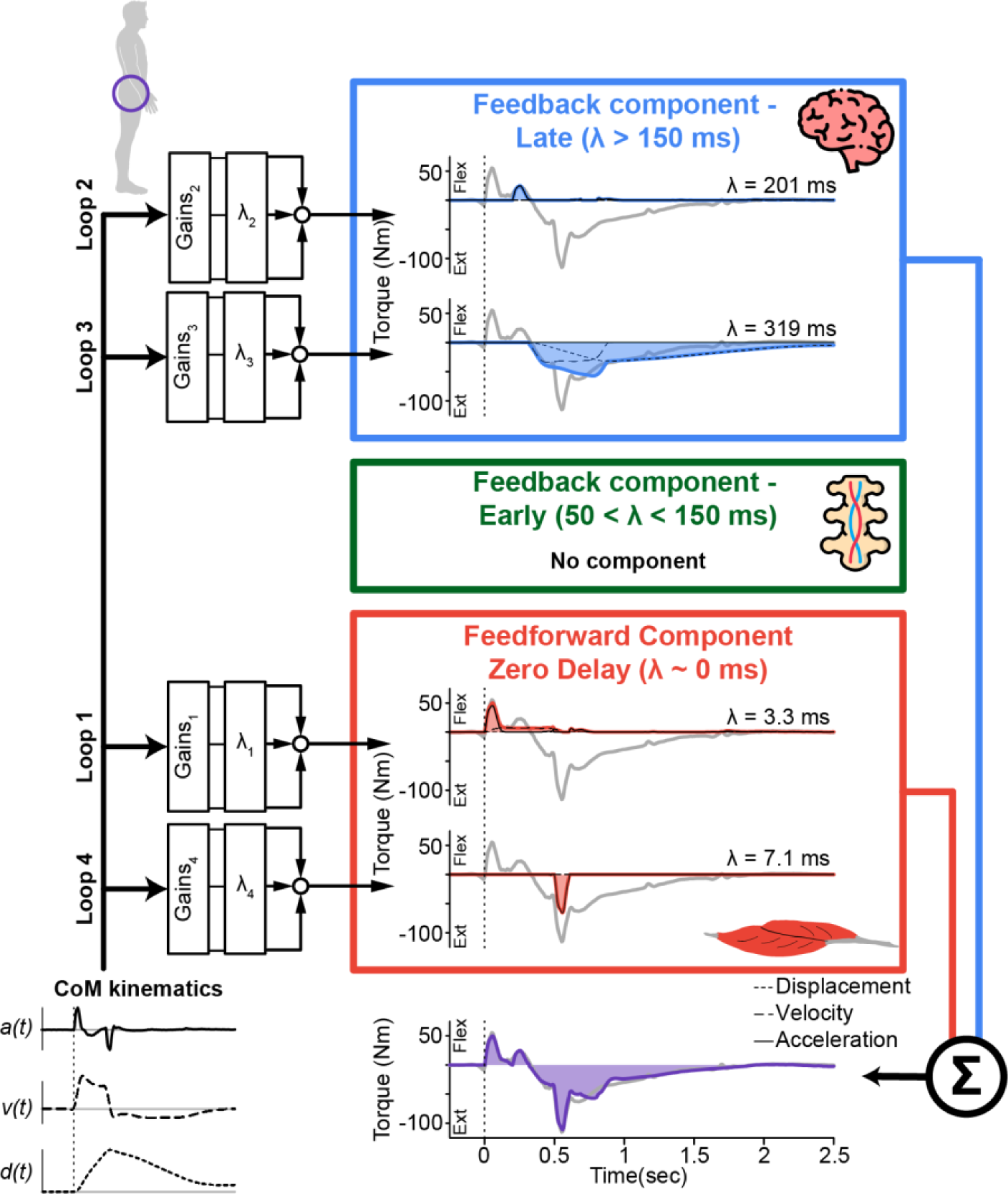
Multi-loop SRM at the hip for a perturbation at 95% of step threshold. At the hip, the balance-correcting torque response is mediated by two feedforward components (red), corresponding to the acceleration and braking of the center of mass, and by two “late” feedback components (blue) with delays longer than 150 ms. The SRM included two loops for any positive change in torque from the baseline and two loops for any negative change in torque from the baseline. The loops are summed, resulting in the overall torque response (purple).

The second hip flexion peak was driven by the acceleration of the CoM at the onset of the ramp perturbation and captured by a second feedback loop with a “late delay” (average *λ_2_* = 160 ± 84 ms across all perturbation magnitudes; Fig 5). Given that this loop was also primarily driven by the acceleration of the CoM as well as the latency and sign of this response (e.g., hip flexion torque), this portion of the response may be elicited by the initial stretch of the hip flexors (e.g., sensory signals from loop 1 drives the sensorimotor response associated with loop 2).

The third SRM loop captures the majority of the hip extension torque response. It was driven by CoM displacement and velocity and had a “late delay” (average *λ_3_* = 329 ± 74 ms across all perturbation magnitudes; Fig 5). A hip extension torque is the expected torque response from posterior chain muscles that will stabilize the body (43).

The final loop captured the peak in the hip extension that was driven by the braking of CoM at the end of the ramp perturbation and occurred at a “zero-delay” loop (average *λ_4_* = 6 ± 9 ms across all perturbation magnitudes; Fig 5).

Across all loops, the delays (λ) did not significantly vary across perturbation magnitudes; however, gains within the first loop did vary (Fig 6). There was a modest, but significant, difference in *K_A1_* during the smallest perturbation (e.g., 12cm) compared to all other perturbations. *K_A1_* was 7% (p=0.004), 8% (p<0.001), and 9% (p<0.001) lower during the 12cm perturbation compared with the perturbation at 75, 85, and 95% of the step threshold, respectively. The difference in the *K_A1_* gain likely indicates a scaling of the feedforward short-range stiffness response with perturbation magnitude. No other gains varied significantly with perturbation magnitude.

**Figure 6.**
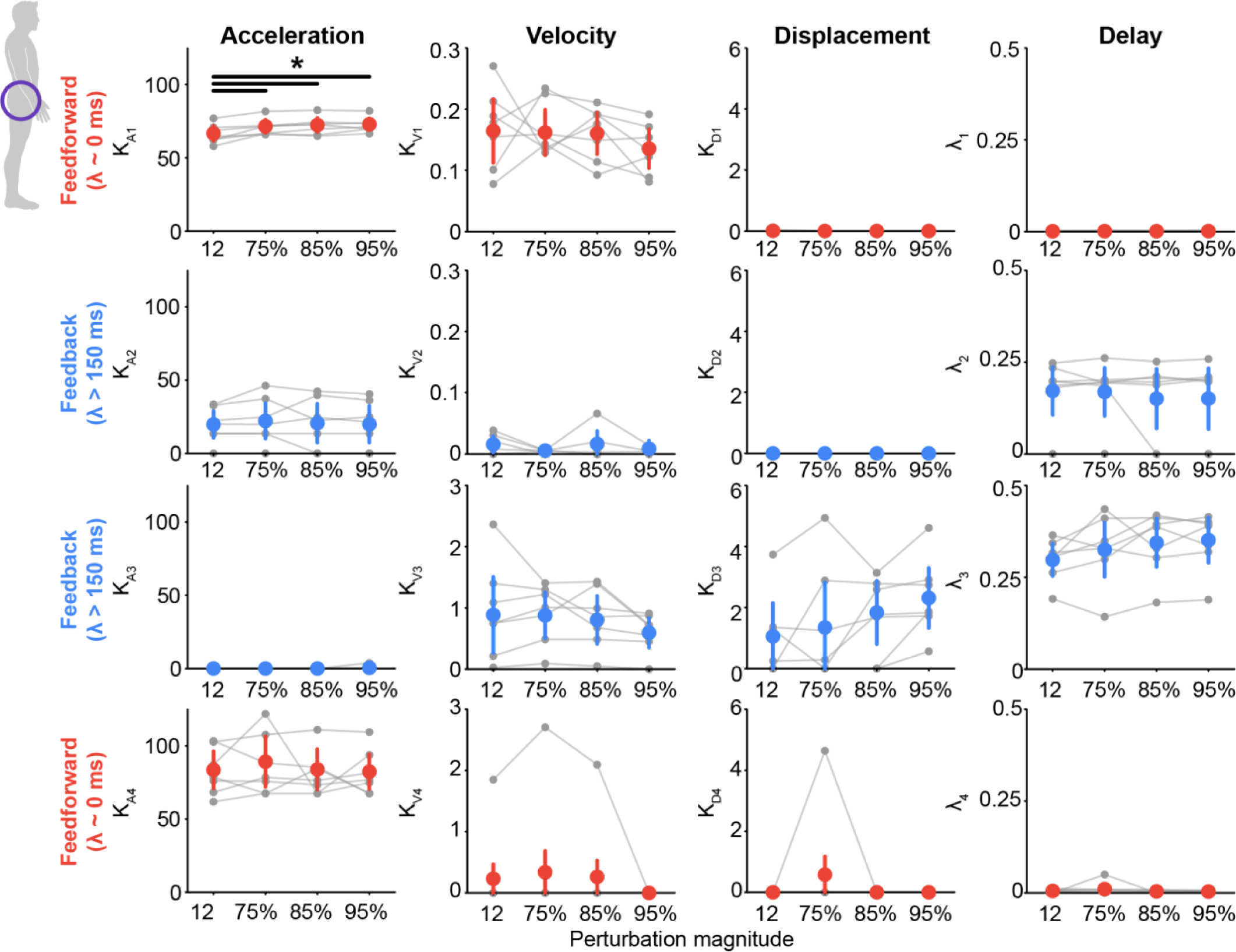
Sensorimotor response model (SRM) gains at the hip for each perturbation magnitude. Each loop was separated into feedforward contribution (red), early feedback contribution (green), or late feedback contribution (blue) based on its delay (λ). *K_Di_*, *K_Vi_*, and *K_Ai_* are the designated SRM gains for CoM displacement, velocity, and acceleration, respectively, while *λ_i_*designates the time delay, *i* represents the *i*th loop. The dots represent the group means and standard deviation, while the gray dots and lines are each participant. The black line and asterisks indicate a significant difference in the SRM gains or time delays across perturbation magnitudes (p < 0.05/6 using Bonferroni corrections for multiple comparisons).

**Knee Response:** The torque response at the knee was captured by three loops, two feedforward loops driven by the acceleration and braking of the CoM, and one feedback loop (Fig 7). The first SRM loop was driven by the acceleration of the CoM at the onset of the ramp perturbation and captured the initial extension torque that occurred at nearly a “zero-delay” (average *λ_1_* = 1 ± 2 ms across all perturbation magnitudes). This response is likely arising from muscle short-range stiffness that is elicited due to the initial flexion of the knee

**Figure 7.**
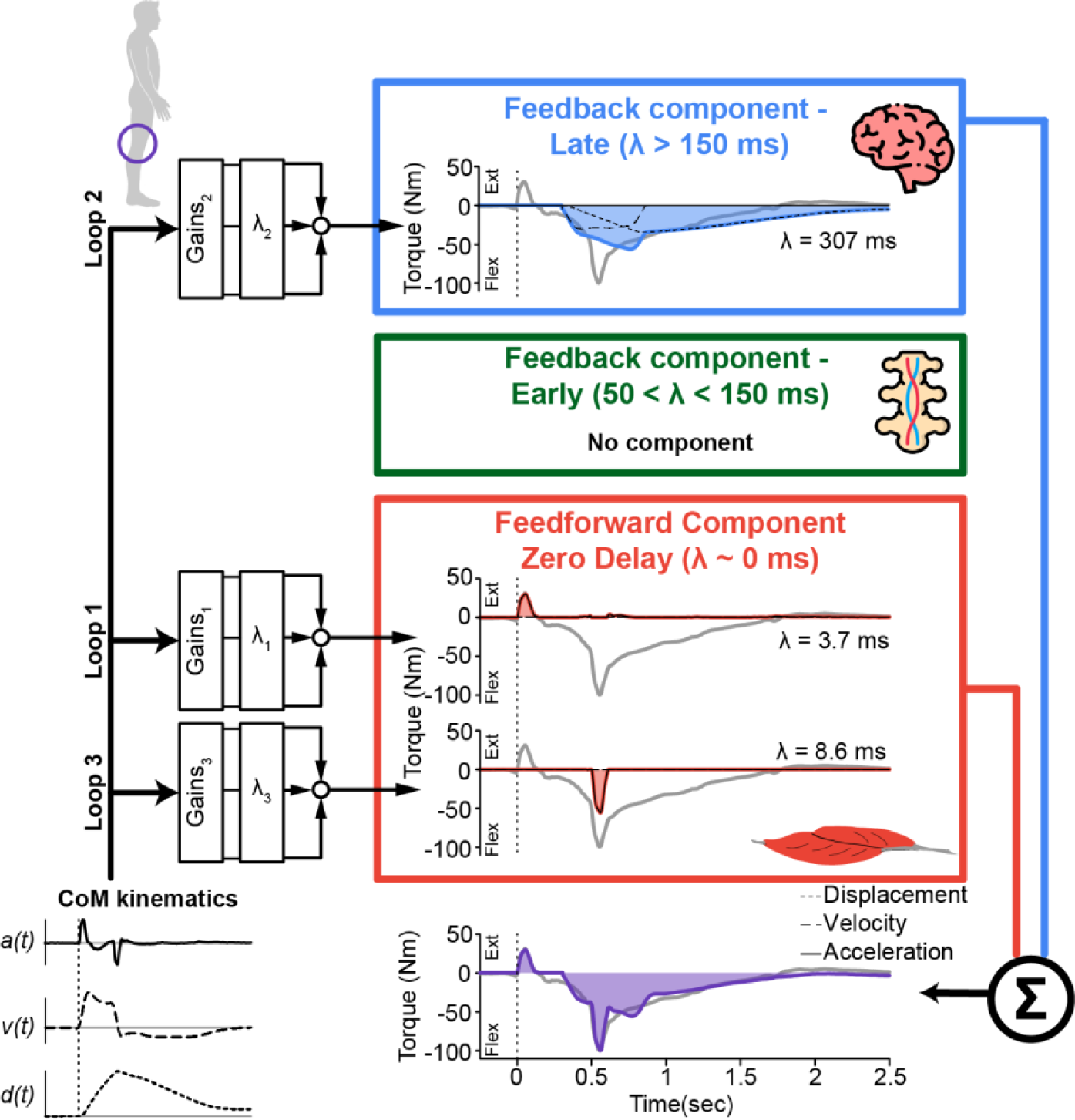
Multi-loop SRM at the knee for a perturbation at 95% of step threshold. At the knee, the balance-correcting torque response is mediated by two feedforward components (red), corresponding to the acceleration and braking of the center of mass, and by one “late” feedback component (blue) with a delay longer than 150 ms. The SRM included one loop for any positive change in torque from the baseline and two loops for any negative change in torque from the baseline. The loops are summed, resulting in the overall torque response (purple).

The second loop captured a majority of the knee flexion torque response. It was driven by CoM displacement and velocity and had a “late delay” (average *λ_2_* = 213 ± 93 ms across all perturbation magnitudes; Fig 7).

The third loop captured the peak in the knee flexions torque that was driven by the braking of CoM at the end of the ramp perturbation, and occurred at a “zero-delay” loop (average *λ_3_* = 4 ± 9 ms across all perturbation magnitudes; Fig 7). It is worth highlighting that the feedforward responses at both the hip and knee may be elicited by the same muscles (e.g., acceleration – rectus femoris; braking – biceps femoris). This is to say that the biarticular muscles that flex the hip also extend the knee, and the biarticular muscles that extend the hip also flex the knee, thus providing a short-range stiffness response at both joints with similar delays.

Across all knee torque loops, the delays (λ) did not significantly vary across perturbation magnitudes; however, the gains did vary within the first loop (Fig 8). There was a modest, but significant decrease in *K_A1_* during the 12cm perturbation compared with the perturbation at 85 and 95% of the step threshold (*12cm vs. 85%*: difference = 10%, p = 0.008, *12cm vs. 95%*: difference = 12%, p = 0.004). It was also significantly lower during the 75% perturbation compared to the perturbation at 95% of step threshold (difference = 4%, p = 0.003). *K_V1_* was significantly higher during the 75% perturbation compared to the perturbation at 95% of step threshold (difference = 41%, p = 0.003). Lastly, *K_D1_* was significantly higher during the 12cm perturbation compared with the perturbation at 75, 85, and 95% of the step threshold (*12cm vs. 75%*: difference = 200%, p < 0.001, *12cm vs. 85%*: difference = 200%, p < 0.001, *12cm vs. 95%*: difference = 200%, p < 0.001). These differences may reflect a scaling of the feedforward short-range stiffness response with perturbation magnitude. No other gains varied significantly with perturbation magnitude.

**Figure 8.**
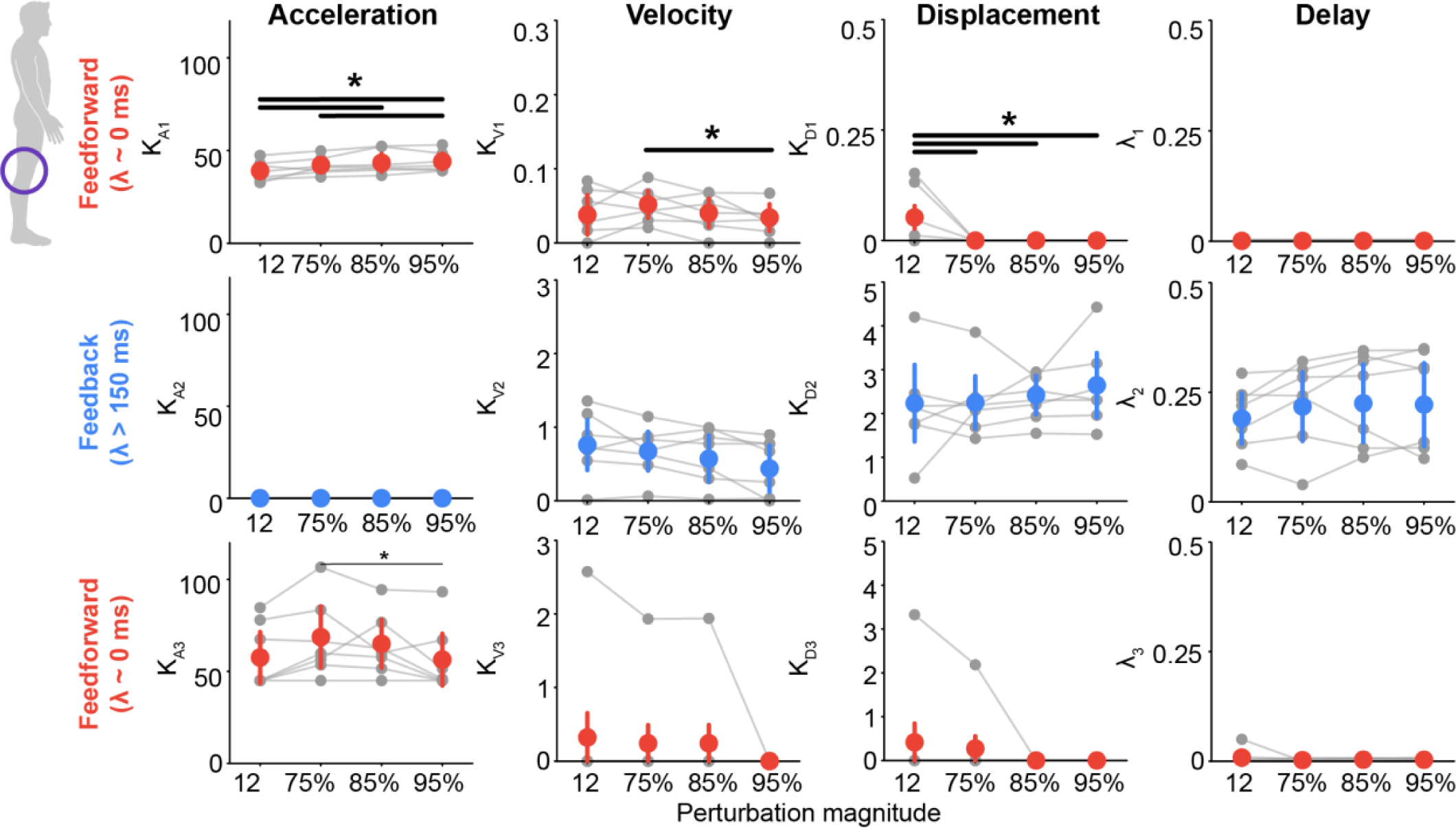
Sensorimotor response model (SRM) gains at the knee for each perturbation magnitude. Each loop was separated into feedforward contribution (red), early feedback contribution (green), or late feedback contribution (blue) based on its delay (λ). *K_Di_*, *K_Vi_*, and *K_Ai_* are the designated SRM gains for COM displacement, velocity, and acceleration, respectively, while *λ_i_* designates the time delay, *i* represents the *i*th loop. The dots represent the group means and standard deviation, while the gray dots and lines are each participant. The black line and asterisks indicate a significant difference in the SRM gains or time delays across perturbation magnitudes (p < 0.05/6 using Bonferroni corrections for multiple comparisons).

**Ankle Response:** In contrast to the hip and knee, the response at the ankle only required feedback contributions, with one “early” feedback loop and two “late” feedback loops (Fig 9). Most notably, there was no “zero-delay” feedforward component in the ankle torque response. This is presumably due to the compliance of the Achilles tendon, which attenuates the short-range stiffness response from the triceps surae.

**Figure 9.**
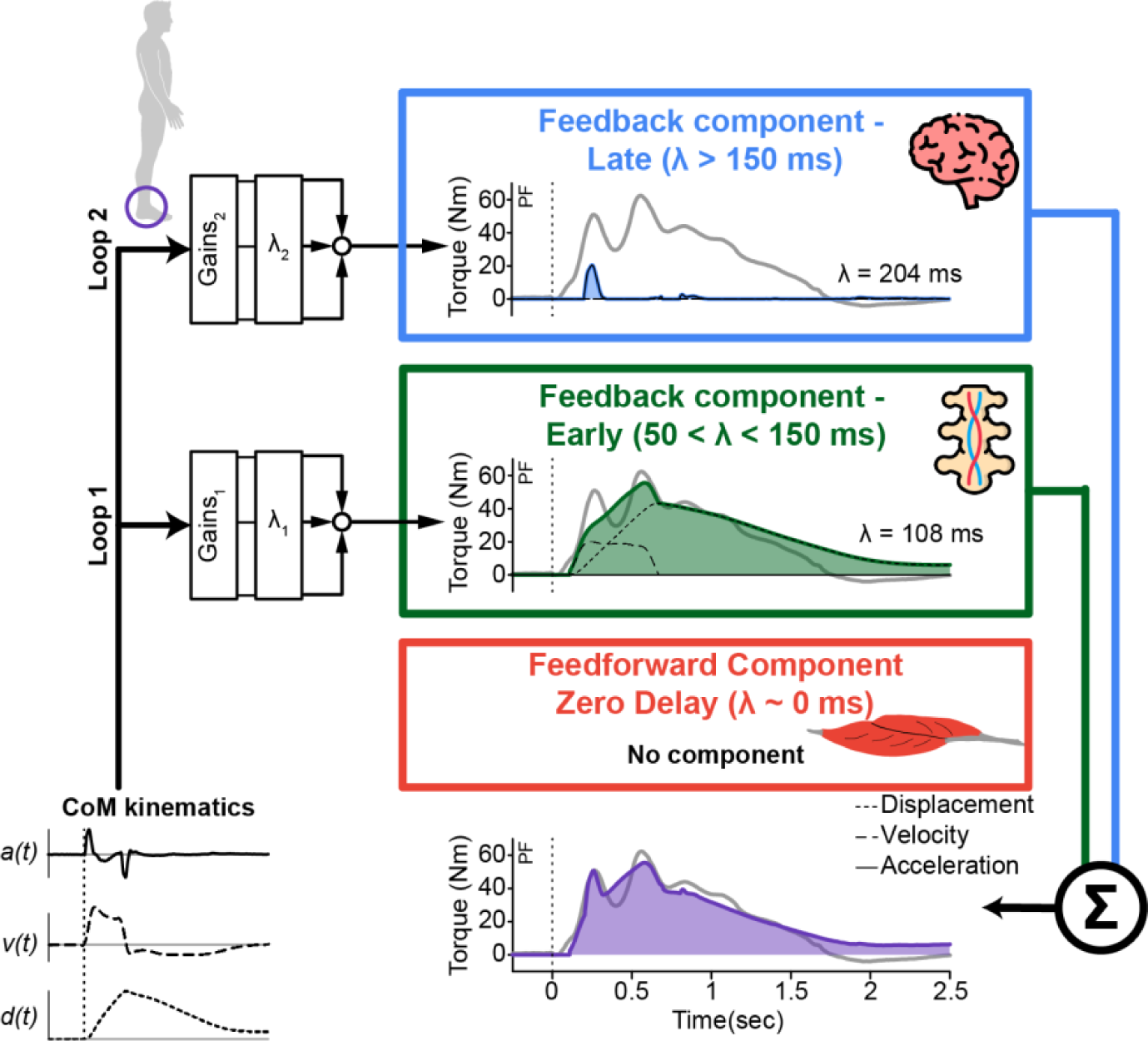
Multi-loop SRM at the ankle for a perturbation at 95% of step threshold. At the ankle, the balance-correcting torque response is mediated by one “early” feedback component that has a delay less than 150 ms and one “late” feedback component with a delay longer than 150 ms. Notably, unlike the hip and knee, there is no feedforward component. The SRM included two loops for any positive change in torque from the baseline and one loop for any negative change in torque from the baseline. The loops are summed, resulting in the overall torque response (purple). Note that in some, but not all participants, a third loop was required; however, it is not shown here.

The first loop captured a majority of the plantarflexion response. It was driven by CoM displacement and velocity and had an “early delay” (average *λ_1_* = 85 ± 23 ms across all perturbation magnitudes; Fig 9). A plantarflexion torque is the expected torque response from posterior chain muscles that will stabilize the body (43).

The second loop captured the first peak in ankle plantarflexion. Interestingly, it was driven by CoM acceleration, but was a “late delay” loop (average *λ_2_* = 190 ± 44 ms across all perturbation magnitudes).

The velocity gain in the first loop (*K_V1_*), as well as the delay of the second loop (*λ_2_*), significantly varied across perturbation magnitudes (Fig 10). *K_V1_* was significantly higher during the 12cm perturbation compared with the perturbation at 95% of the step threshold (difference = 73%; p = 0.005). It was also higher during 75% compared with 95% of the step threshold (difference = 51%; p = 0.004). As gains do not typically decrease as perturbation magnitude increases, this may represent a saturation of CoM velocity in the ankle response. There was a modest, but significant difference in *λ_2_* during the 75% compared with 95% of the step threshold perturbations (difference = 4%, p = 0.008).

**Figure 10.**
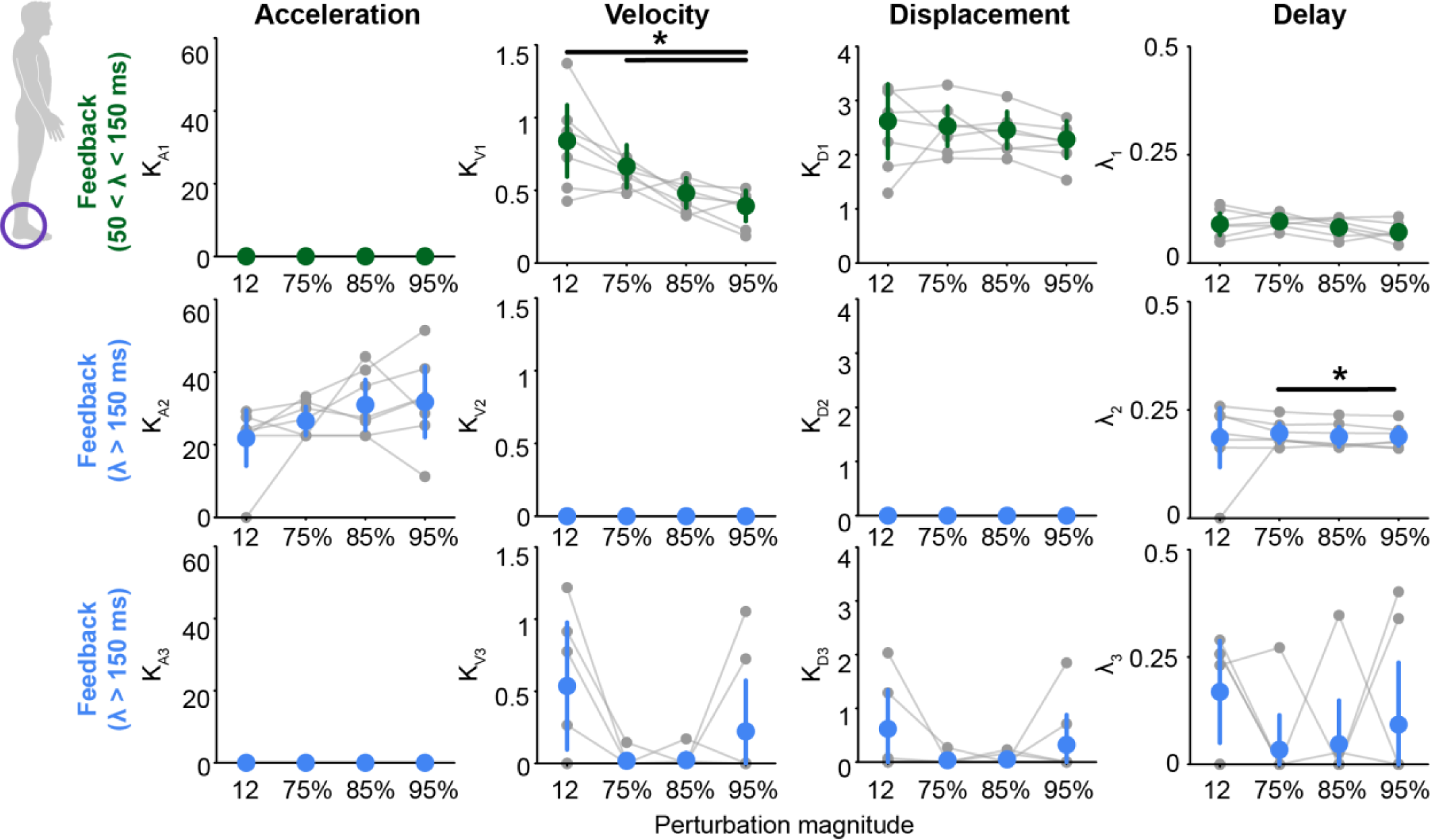
Sensorimotor response model (SRM) gains at the ankle for each perturbation magnitude. Each loop was separated into feedforward contribution (red), early feedback contribution (green), or late feedback contribution (blue) based on its delay (λ). *K_Di_*, *K_Vi_*, and *K_Ai_* are the designated SRM gains for COM displacement, velocity, and acceleration, respectively, while *λ_i_*designates the time delay, *i* represents the *i*th loop. The dots represent the group means and standard deviation, while the gray dots and lines are each participant. The black line and asterisks indicate a significant difference in the SRM gains or time delays across perturbation magnitudes (p < 0.05/6 using Bonferroni corrections for multiple comparisons).

## DISCUSSION

Our work provides novel insight into how neurally-mediated feedforward and feedback pathways contribute to the overall multi-joint torque response, supporting our secondary hypothesis that a torque-SRM could differentiate feedforward and feedback contributions to the torque response at each joint. Our results also indicate that the reactive torque response at the hip, knee, and ankle can be robustly described by sensorimotor feedback of center of mass kinematics, supporting the established hypothesis that the nervous system uses task-level variables to drive the coordinated multi-joint response (23, 29, 31, 32). Interestingly, the pathways contributing to the overall response varied at each joint, indicating that while a task-level variable, CoM kinematics, drives the torque response, the response is joint-specific. Variation between joints may be attributed to differences in musculotendon mechanical properties between proximal and distal joints, as well as differences in the elicited sensory feedback pathways. For example, at the hip and knee, we found a feedforward torque response to the acceleration and braking of the CoM, as well as “late” feedback responses. In contrast, at the ankle, we only observed feedback contributions, with one being an “early” response and the others being “late” feedback responses. The lack of a feedforward contribution at the ankle may be driven by the compliance of the Achilles tendon, which attenuates the intrinsic mechanical response from muscle short-range stiffness. Differentiating the feedforward and feedback contributions at each joint can aid in our understanding of how each joint contributes to the balance-correcting response and how the response can be modulated. It can also aid in identifying disrupted pathways that result in impaired balance in older adults or those with neuromuscular injuries or diseases. Lastly, the ability to mimic the physiological balance-correcting torque response may aid in developing legged robots and wearable robotic devices that can withstand and help the user withstand postural perturbations, respectively.

### Feedforward and feedback contributions to the reactive torque differ across joints

Distal tendons that are more compliant may attenuate the muscle short-range stiffness response, leading to the lack of feedforward response at the ankle. At the hip and knee, we observed “zero-delay” feedforward loops, while a “zero-delay” loop was not present at the ankle (Fig 5, 7, & 9). The “zero-delay” feedforward torque presumably arises from the intrinsic mechanical properties of the musculoskeletal system, including muscle short-range stiffness. The muscle short-range stiffness response would be elicited immediately following the perturbation and prior to muscle activation, as we found (Fig 2). Additionally, the magnitude of the short-range stiffness response is sensitive to the amplitude of the stretch within the muscle (13). Due to the serial connection between muscle and tendon, the stretch that occurs within each muscle during the imposed perturbations will be dependent upon the compliance of the tendon to which it is attached, with more stretch occurring within the tendon when it is more compliant than the muscle. Our results suggest that, at the ankle during postural conditions, since the tendon is less stiff than the muscle at nearly all levels of muscle activation (44-46), a majority of the perturbation-related stretch occurs within the tendon, attenuating the short-range stiffness response within the muscle and no “zero-delay” feedforward component at the ankle. In contrast, at the hip and knee, which are thought to have stiffer tendons (47), most of the imposed stretch occurs within the muscle, resulting in the “zero-delay” feedforward torque. This result is supported by prior musculoskeletal modeling work that found that including muscle short-range stiffness within the model resulted in hip and knee torques being generated prior to muscle activation but not ankle torques (30). We note the small rise in ankle torque at the time of the perturbation that is unaccounted for in our current model. However, even if we implemented a “zero-delay” loop, it was unable to capture this response. This could be due to non-linear musculotendon mechanics that our linear model was unable to capture.

Differences in tendon stiffness between the proximal and distal joint may also impact the efficacy of using feedforward control, and the lack of a “zero-delay” feedforward torque response at the ankle has important implications for balance control. Feedforward modulation of muscle activity or co-contraction is thought to increase muscle stiffness, thereby increasing the resultant feedforward torque, which can improve postural stability by providing greater instantaneous resistance to unexpected perturbations (9, 10). However, our results suggest that feedforward modulation at the ankle may be an ineffective way to improve postural stability. Due to the compliance of the Achilles tendon relative to that of the triceps surae (44-46), feedforward increases in muscle activation would result in a minimal increase in the resultant ankle torque that arises from muscle short-range stiffness. This is in agreement with previous findings that ankle stiffness is insufficient to maintain postural stability (20, 22). In contrast, since the tendons at the hip and knee are likely less compliant than the Achilles tendon (47), increasing muscle activation or co-contraction at the hip or knee may be an effective way to increase the feedforward torque response.

While a global change in CoM drives the feedback response at each joint, our results suggest that different neural mechanisms may modulate the feedback responses across the hip, knee, and ankle. Through the use of the torque-SRM, we separated the feedback pathways into “early” and “late” components based on the time delays of each loop (Fig 5, 7, & 9). These results vary from previous findings where the same CoM kinematics transformation, with a single delay, could predict the coordinated muscle activity across different joints (48). However, this model only had a single delay consistent with the sub-cortical response. It was recently observed that implementing a parallel loop EMG-SRM to fit medial gastrocnemius activation could capture both cortical and sub-cortical contributions, significantly improving the overall fit (28). If reactive EMG signals at the ankle arise from both cortical and sub-cortical pathways, so would the resultant ankle torque, as we found in our study. Interestingly, the same cortical and sub-cortical pathways do not appear to be modulating the resultant torques across joints. For example, at the hip, one feedback pathway is likely transcortical (average delay of 160 ± 84 ms) while the other may be a voluntary response (average delay of 329 ± 74 ms; Fig 6) (28, 49). In contrast, at the ankle, there was an “early” loop (*λ_1_* = 85 ± 23 ms) that was likely a spinal or brainstem mediated pathway, while the “late” loop is likely a transcortical pathway (*λ_2_* = 190 ± 44 ms) (1, 28, 49). While we can speculate on the neural origin of each feedback pathway, definitively identifying the sensory feedback pathway for each loop was outside the scope of this study and requires future investigation.

### Limitations

One limitation of the current study is that only a single perturbation direction (backward support surface translations) was tested. Due to differences in musculotendon architecture, the feedforward contribution may vary with perturbation direction. The torque-SRM’s ability to predict joint torque has only ever been evaluated in the sagittal plane (29). Thus, it is unclear if the torque-SRM can predict frontal plane joint torques. While the EMG-SRM has accurately predicted reactive muscle activations during frontal plane perturbations (24), future work should include testing the efficacy of the torque-SRM at capturing frontal plane torques.

The perturbations tested were also large (at 95% of the step threshold). The response to these perturbations required both a hip and ankle strategy, even during the smallest applied perturbation (12cm). It remains to be seen if the torque-SRM introduced in this study is robust during smaller perturbations that mainly require an ankle strategy. Additionally, the participants were instructed to maintain a foot-in-place balance response, even to the largest applied perturbations. Since the perturbations were close to the participant’s step threshold, it is possible that the natural response would have been to take a step. Additional testing is required to determine if the torque-SRM can accurately predict the reactive joint torques when a step is taken.

While global changes in CoM kinematics drive the feedback neural responses, it is possible that CoM kinematics are not the physiological driver of the feedforward muscle short-range stiffness response. We found that global changes in CoM kinematics modulate the balance-correcting torque response. This builds on prior work that has demonstrated that reactive muscle activations are driven by global changes (e.g., CoM) rather than local, joint or muscle level changes (27, 50-52). Since CoM is an abstract, global variable, within this framework, sensory information from proprioceptive, visual, and vestibular systems are integrated to generate an appropriate corrective torque (53), presumably within the brain stem or cortex. However, muscle spindle activity, which contributes to the proprioceptive response and is related to muscle short-range stiffness, is activated in response to the initial stretch acceleration (or the first time derivative of force dF/dt) within the muscle (e.g., a local signal) (54). One of the biggest limitations when exploring if global versus local signals drive the feedforward muscle short-range stiffness response is that we do not have a good measure of angular acceleration that could be driving this response. Thus, it is possible that joint angular accelerations drive the feedforward response, and CoM acceleration is a proxy measure.

### Future implications and conclusions

The ability to differentiate the feedforward and feedback contributions, as well as the different feedback pathways that are contributing to the overall multi-joint torque response, may provide a framework for determining mechanisms underlying the impaired control of balance in aging, injury, or neuromuscular pathology. For example, older adults have decreased Achilles tendon stiffness (55-57), and an increase in the delay of the feedback pathways (58-60). This methodology could differentiate the impact of these changes on the overall balance-correcting response. Once the deficit is identified, targeted training at the source of the impairment can be developed. This is critical since training or treatment targeted at neural deficits (e.g., sensory feedback delays) will vary from training targeted at biomechanical deficits (e.g., tendon stiffness). This same framework could be used to identify specific deficits in individuals with Parkinson’s disease, older adults with mild cognitive impairment, stroke survivors, or other neuromuscular injuries and diseases.

Our framework may also simplify the control of legged bi-pedal robots, and lower-limb prostheses and exoskeletons. Our framework uses CoM kinematics, a single control signal, to predict the torque response at the ankle, knee, and hip. This one-to-many mapping, rather than the one-to-one mapping currently employed, could simplify the control of these devices. Moreover, using a physiologically-inspired control scheme, where the controller mimics the biological feedforward and feedback responses to postural perturbations, may also improve the embodiment of devices. The principles of embodiment suggest that robotic devices should coordinate with the human’s natural response, such that the nervous system can model the controller of the robotic device (61). Since a torque-SRM control scheme would be based on the nervous system’s response, very little learning might be required for the human to model the controller. This could ultimately improve device acceptance and usage in the real world.

## DATA AVAILABLITY

The data from the current study are available from the corresponding author upon reasonable request.

## GRANTS

This publication was supported by grant number 2127509 from the NSF and American Society for Engineering Education, National Institutes of Health grants F32 AG063460, R01 HD046922, R01 HD090642, and McCamish Parkinson’s Disease Innovation Program. Its contents are solely the responsibility of the authors and do not necessarily represent the official views of the National Science Foundation, American Society for Engineering Education, National Institutes of Health, or McCamish Foundation.

## AUTHOR CONTRIBUTIONS

G.M., O.N.B., K.L.J., and L.H.T. conceived and designed research; G.M. and O.N.B. performed experiments; K.L.J analyzed data, K.L.J., G.S.S., and L.H.T interpreted results of experiments, K.L.J prepared figures and drafted manuscript, K.L.J., G.M., O.N.B., G.S.S., and L.H.T. approved final version of manuscript

